# An efficient algorithm for Nash equilibria when harvesting interacting living resources

**DOI:** 10.1101/2023.01.25.525330

**Authors:** T. J. Del Santo O’Neill, A. G. Rossberg, R. B. Thorpe

## Abstract

Natural ecological communities exhibit complex mixtures of interspecific biological interactions, which makes finding optimal yet sustainable exploitation rates challenging. Most fisheries management advice is at present based on applying the Maximum Sustainable Yield (MSY) target to each species in a community by modelling it as if it was a monoculture. Such application of single-species MSY policies to strongly interacting populations can result in tragic overexploitation. However, the idea of “maximising the yield from each species separately” can be extended to take into account species interactions using a multispecies or ecosystem model and determining a Nash Equilibrium (NE), where the yields of each species taken in isolation are simultaneously maximised. Here we present ‘`nash`’, an `R` package that streamlines the computation of NE reference points for fisheries and other systems represented by a user-defined multispecies or ecosystem model. We present two real-world fisheries management applications alongside performance benchmarks. Satisfactorily search results are shown across models with an approximate factor 15 increase in performance when compared to the expensive round-robbing sequential optimisation algorithms used by other authors in the literature. We believe that the `nash` package can play an instrumental role in fully implementing ecosystem-based management policies worldwide.

**Open Research statement:** This submission uses novel code, which is provided, per our requirements, in an external repository to be evaluated during the peer review process via the following link https://github.com/ThomasDelSantoONeill/nash

## 1 Introduction

Public environmental policies conceptualise the natural world as a stock of materials that yield flows of goods and/or ecosystem services^1^ (*e*.*g*. food production from fisheries) to society (Costanza et al., 1997; Daily, 1977; Millennium Ecosystem Assessment, 2005; Costanza et al., 2017). Use of these services is considered *sustainable* if, at a minimum, it can indefinitely be continued at equal levels into the future. With this notion of sustainability (although ‘stronger’ interpretations exist *cf* Daly, 1995; Figge, 2005; Kuhlman and Farrington, 2010; Rossberg et al., 2017), when decision makers seek to attain the highest possible sustainable flow rates, this objective is called *Maximum Sustainable Yield* (MSY).

Whilst the MSY concept is applicable to a wide range of natural systems, we here focus on the important example of fisheries. The current practise of fisheries management is still largely based on a *stock-by-stock* ^2^ or ‘single-species’ (SS) MSY approach; whereby the indirect effects of fishing through biological interactions and the intrinsic dependencies with other abiotic ecosystem components are not explicitly dealt with (Skern-Mauritzen et al., 2016; Marshall et al., 2018; Howell et al., 2021). Indeed, previous contributors to *Ecological Applications* have argued that sustainability is “more akin to the simultaneous [sustained yield]^3^ of many interrelated populations in an ecosystem” (Goodland and Daly, 1996) and that the “goal of sustainable yield of single species in a fishery is a fundamental mistake” (Pitcher, 2001), asking “what do we mean by MSY when the interactions between species are taken into account?” (Andersen et al., 2015). Here we reconcile these concerns by presenting a novel application that (i) implements a specific formal interpretation of MSY that aligns with current policy objectives whilst considering interactions and (ii) streamlines its computation for any model that, at the very least, acknowledges trophic interactions; that is, any multispecies model in the sense of Link’s (2002) “gradient” to fisheries management.

The MSY concept was first devised in the context of a fishery in the 1930s (Russell, 1931; Hjort et al., 1933; Graham, 1935) and following the 1982 United Nations Convention on the Law of the Sea (UNCLOS)^4^, it was enshrined as the nominal objective in the legislation of different jurisdictions worldwide; examples are the U.S. Magnuson-Stevens Fishery Conservation and Management Act (MSA)^5^, the EU Common Fisheries Policy (CFP)^6^ and UK’s Fisheries Act^7^.

Having been subjected to scrutiny since its inception, the MSY objective *per se* has remain largely unchanged except for its occasional reinterpretation as an upper harvesting ‘limit’ (Mace, 2001; Quinn and Collie, 2005). Our application offers the option to set lower limits on stock sizes and so upper limits on exploitation rates to implement such “constraints of biodiversity conservation” (*sensu* Matsuda and Abrams, 2006; see section 2.4 dor details). Scepticism over MSY arose (*e*.*g*. Gordon, 1954; Larkin, 1977; Sissenwine, 1978; Cunningham, 1981; Yodzis, 1994; Punt and Smith, 2001) from a view of the resource unit as part of a larger, highly coupled, social-ecological system (*sensu* Ostrom, 2009). As a result, there has been a shift towards more holistic management paradigms, such as the Ecosystem Approach to Fisheries Management (EAFM), where some ecosystem considerations are incorporated (though seldom quantitatively) into SS stock assessment models (Skern-Mauritzen et al., 2016; Marshall et al., 2018; Pepin et al., 2022). However, EAFM does not always mean that biological interactions are fully represented in models (Link, 2010; Link and Browman, 2014). Advice by the International Council for the Exploration of the Sea (ICES), for example, considers interactions by rescaling natural mortality rates based on information derived from multispecies models (ICES, 2022a; Plagányi et al., 2022). In Europe, EAFM has been operationalised through the Marine Strategy Framework Directive (MSFD)^8^ and several multiannual plans^9^.

It is known that the sustainable SS yield maxima for different species correspond to different ecosystem states and are therefore not simultaneously attainable (*e*.*g*. Pope, 1976; May et al., 1979; Beddington and May, 1980; Walters et al., 2005; Mackinson et al., 2009; Legović and Geček, 2010; Fogarty, 2014; Moffitt et al., 2016; Link, 2018). The issues become most apparent when different stocks are exploited by different agents. For instance, in a predator-prey system maximising the yield of the predator requires its prey to be unexploited, whereas by harvesting the predator, the yield of the prey can be maximised. A conflict arises when two managers, each targeting either the predator or the prey, seek to maximise their individual yield whilst disregarding the other. Naturally, such conflicts increase as the number of species and/or interest groups become larger (Cochrane, 1999).

Addressing the potential shortcomings of SS-MSY policy objectives, Farcas and Rossberg (2016) investigated the effectiveness of several alternative multispecies translations of SS-MSY. Based on considerations of yields attained, transparency of negotiations, impacts on biodiversity, and continuity with the existing European framework, they recommended to “maximise the yield from each stock separately” in the sense of a Nash (1951) equilibrium (NE) with respect to fishing mortality rates. This objective, abbreviated NE-MSY below, is attained if each targeted stock of a wild mixture is harvested at such rates that no deviation from this rate can increase the long-term yield from that species. If each stock was fished by an independently but rationally managed fleet, this Nash equilibrium would be the natural long-term outcome.

Several authors have since demonstrated the advantages of this form of NE-MSY. For instance, Norrström et al. (2017) demonstrated the existence of such a solution in the Baltic Sea fishery and Thorpe et al. (2017), Thorpe and De Oliveira (2019), Thorpe (2019) and Spence et al. (2020) concluded that harvesting close to or below NE-MSY generates the greatest collective long-term benefits in the North Sea fishery; in terms of yield weight, economic revenue and risk of stock collapse. It should be noted that NE-MSY is not the only option to simultaneously maximise the yield from each stock (Farcas and Rossberg, 2016). Compared to other options, it is however, conceptually similar to SS-MSY as defined in public policy and thus expected to be more easily accepted by stakeholders.

Despite these well-established strengths of the concept, an efficient algorithm for the computation of NE-MSY for complex ecological food web models remains elusive. The present work fills this gap by implementing an efficient algorithm to calculate NE-MSY in ‘`nash`’, a new `R` package (R Core Team, 2020) which we designed to streamline the computation of NE-MSY objectives for any abstract models where, at a minimum, the targeted species biologically interact with one another. We commence with a thorough description of `nash`’s internal machinery and then provide installation instructions alongside three working example applications and performance benchmarks.

## 2 Materials and Methods

### 2.1 Assumptions

The package `nash` is designed to find NE in the context of ecosystem management. The algorithms implemented in `nash` assume that users have developed an ecosystem model with *S* harvested compartments, typically represented by population biomass variables. In the simplest case, population dynamics are given by a system of autonomous ordinary differential equations of the general form

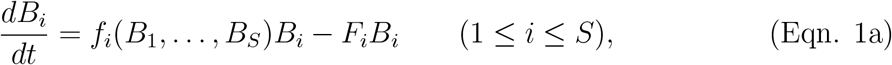

with non-negative biomass or state variables *B*_*i*_ (*i* = 1, …, *S*), and *f*_*i*_(*B*_1_, …, *B*_*S*_) specifying the momentary population growth (or decay) rate of compartment *i* in the absence of exploitation. The latter may depend on the biomass of *i* and of all other compartments in the model. The last term in equation (1a) describes removal (*e*.*g*. harvesting and/or culling) of biomass from compartment *i* at a *harvesting rate F*_*i*_. Note that *F*_*i*_ may be numerically different from the fishing mortality rate if fishing is size or age specific (Worm et al., 2009; Shephard et al., 2012; Fung et al., 2013; Fung, 2013).

Equation (1a) can be reformulated in matrix-vector notation as

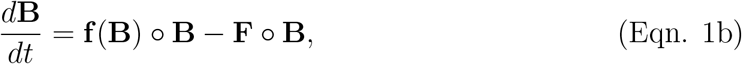

with *°* denoting the entry-wise or Hadamard product. For given **F**, the user-specified model outputs the yields at the stable equilibrium **Y** = **F** *°* **B**^*^ = **F** *°* **B**^*^(**F**) of equation (1b), assumed to be unique.

However, `nash` can also handle situations where the user-defined model contains other variables that do not correspond to harvested population sizes (*e*.*g*. unharvested populations or variables describing population structure), where different components of a population (*e*.*g*. different age classes) are fished at different rates, and situations where the model steady state is not a fixed point equilibrium but instead, *e*.*g*. a limit cycle.

In this case the user-supplied model should output as the vector **Y** the steady-state averages of the total yields from the harvested populations. For each combintion of input parameters **F** and outputs **Y**, one can define *nominal effective equilibirum biomasses* **B**^*^as 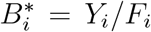. A situation similar to the equilibrium condition of equation (1b) is then satisfied if **f** is *redefined* as follows: for a given vector **B**^*^ the value of **f** (**B**^*^) is given by the value of **F** chosen such that, in simulations of the ecosystem model, the steady state averages of the biomasses become exactly **B**^*^. By this definition, 0 = **f** (**B**^*^) *°* **B**^*^ − **F** *°* **B**^*^, which is analogous to the equilibrium condition of equation (1b).

### 2.2 Theoretical Framework

General ecological models following equation (1) can be approximated near equilibria by adequately calibrating generalised Lotka-Volterra models (MacArthur, 1970; Gilpin and Justice, 1973; Kirkwood, 1982). This is achieved by linearising *f*_*i*_(*B*_1_, …, *B*_*S*_) by a first order Taylor series expansion around the equilibrium **B**^*^:

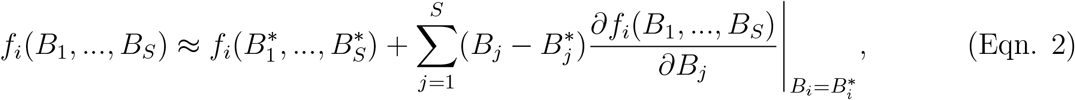

In this expression, the term 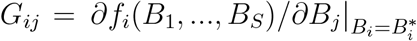 provides the elements of the so-called *interaction matrix* **G** (Berlow et al., 2004; Novak et al., 2016). *G*_*ij*_ quantifies the direct effects per unit of population biomass that species *j* has on the growth rate of species *i*. Therefore, the matrix **G** describes the combined effects of, *e*.*g*., competitive, exploitative, or mutualistic interactions.

Defining the constants *r*_*i*_ as being equal to the terms 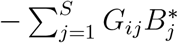 in equation (2), and putting equation (2) into equation (1) yields the Lotka-Volterra approximation of the dynamics in the neighbourhood of **B**^*^ as

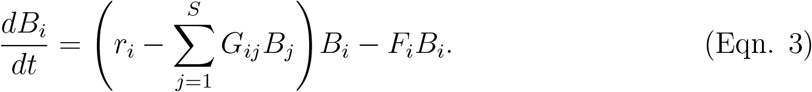

In the fisheries literature, this model is referred to as the multispecies extension of Schaefer’s (1954) *surplus production model* (Pope, 1976) or the Lotka-Volterra model (Lotka, 1925; Volterra, 1926).

For any parameterisation of equation (3) where the populations in all *S* compartments coexist 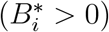, equation (3) implies (in vector-matrix notation)

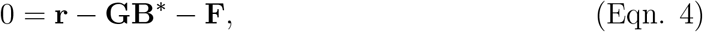

which can be solved for the equilibrium biomasses as

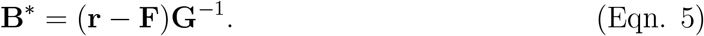

### 2.3 Nash Equilibrium

Formally, NE-MSY as defined above corresponds to yields and harvesting rates satisfying the condition

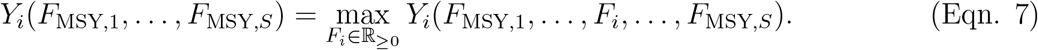

For a sufficiently smooth dependence of yields on harvesting rates (given in almost all ecological situations), maximising the long-term yield of any species *i* is analogous to finding the point at which the first derivative of yield with respect to harvesting rates is zero. For the Lotka-Volterra model equation (3), a NE-MSY is therefore attained if

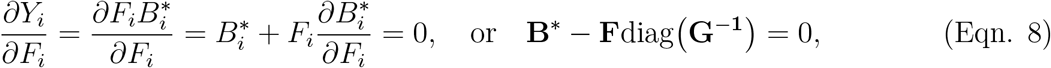

where we used that, by equation (5), 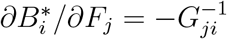. In equation (8) diag(**G**^−1^) denotes the diagonal matrix formed by the elements on the diagonal of **G**^−1^. This result allows us to derive closed formulas for NE-MSY in Lotka-Volterra models.

Specifically, equation (8) can be solved for **F** as

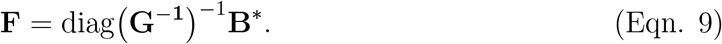

Plugging this into equation (5) and solving for **B**^*^ = **B**_Nash_ gives

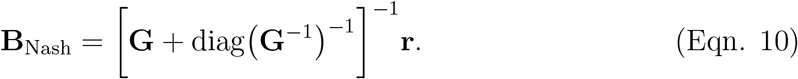

The NE harvesting rates **F** = **F**_Nash_ are then computed by combining equation (9) and equation (10):

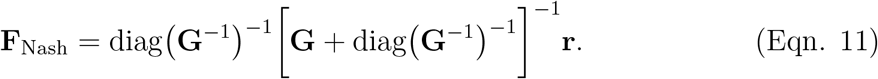

These analytic results are at the heart of the algorithm implemented in `nash`.

### 2.4 Implementing Conservation Constraints

It is conceivable that in some cases harvesting at the Nash equilibrium might drive one or more species to extinction. Even if this does not happen, some species might be placed under unacceptable risk levels of stock collapse. Therefore, the `nash` package includes an option that allows the user to explicitly incorporate conservation constraints (*sensu* Matsuda and Abrams, 2006), specifying a conservation biomass threshold *B*_cons,*i*_ below which stock sizes are not permitted to fall. As a user guideline we note that in Europe, for instance, the International Council for the Exploration of the Sea (ICES) establishes “deterministic biomass limits below which a stock is considered to have reduced reproductive capacity” (ICES, 2022b). Although its computation is a matter of current debate (*cf* ICES, 2022c), a choice common in the literature is to set *B*_lim_ as a fraction (*e*.*g*. 10-25%) of the unfished biomass *B*_0_. Similarly, Minimum Stock Size Thresholds (MSSTs) are set for stocks within the U.S. MSA jurisdiction typically taking the largest biomass value between (i) one-half the biomass that equates to MSY, or (ii) the minimum biomass at which rebuilding to MSY occurs within 10 years if harvested, utmost, at rates associated with MSY (for details see Published Rule 63 FR 24212 of May 1, 1998, by the National Marine Fisheries Service at https://www.federalregister.gov/d/98-11471).

We implement these constraints as follows. If some *B*_Nash,*i*_, computed as above, fall below *B*_cons,*i*_, the species with the lowest *B*_Nash,*i*_*/B*_cons,*i*_ ratio is identified and its target biomass set to *B*_cons,*i*_. Subsequently, eliminating species *i* from equation (3) and re-evaluating equation (10) leads to a new estimation of **B**_Nash_. This procedure is repeated until *B*_Nash,*i*_ *> B*_cons,*i*_ for all remaining species. Finally, equation (9) is evaluated with *B*^*^ replaced by *B*_cons,*i*_ where constraints apply and by *B*_Nash,*i*_ otherwise. By default, all *B*_cons,*i*_ are set to 0 in `nash`, so no conservation constraints are applied.

### 2.5 The nash Function

The function `nash` included in the `nash` package was built with a user interface similar to that of the general-purpose optimisation function `optim` included in `R`. Data structures of inputs and outputs mirrors `optim` as far as feasible.

At a minimum, the `nash` function requires as inputs (i) initial ‘`par`’ values of the *S* parameter that will be optimised (*i*.*e*. the harvesting rates), and (ii) the *S*-valued objective function ‘`fn`’ for which NE-MSY is to be found. Specifically, `fn` takes an atomic vector of real double-precision harvesting rates values as input and outputs an atomic vector of real double-precision long-term yields. Typically, running `fn` will require simulation of a user-defined ecosystem model for given harvesting rates until a steady state is reached.

Running `nash` returns an atomic vector of the ‘list’ type containing (i) the NE harvesting rates in the same format as the `par` input, (ii) the corresponding yields (‘`value`’), (iii) the number of times (‘`counts`’) that `fn` was evaluated, and (iv) an atomic vector of the ‘character’ type indicating successfulness (or not) of convergence.

As an optional argument to `nash`, the user can specify whether to utilise the ‘`LV`’ or ‘`round-robin`’ method to compute NE harvesting rates. The former is described below, whilst the latter was implemented for benchmarking purposes to replicate the algorithm employed by, *e*.*g*., Norrström et al. (2017), Thorpe et al. (2017) or Spence et al. (2020); that is, round-robbing sequential optimisation of each *F*_*i*_ until convergence is reached (see Algorithm 2 for details).

#### 2.5.1 The LV Method

The `LV` method iteratively approximates the model ecosystem by an LV model (section 2.2) and computes NE-MSY based on equation (10) and equation (11) until a convergence thresholds is satisfied, while taking conservation constraints (section 2.4) into account. Details are given in Algorithm 1.

The approximation of the inverse of the **G** matrix in equation (5) is obtained in `nash` through the following 2^nd^ order central finite-difference scheme

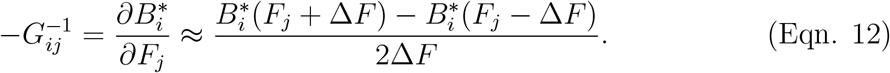

Selecting a small value of the step size Δ*F* results in a more precise estimation of **G**. According to Collie et al. (2003) and Pope et al. (2019), Δ*F* = 0.1 seems a reasonable value (assuming *F* is measured in units of year^−1^) to balance the effects of different sources of numerical error (Higham, 2002; Press et al., 2007). Although Δ*F* is set to 0.1 within `nash` by default, this argument can be user-specified.

### 2.6 Benchmark Models

To demonstrate the use of `nash`, we applied it two three models which are, in order of increasing complexity: (i) Taylor and Crizer’s (2005) modified two-species competitive Lotka-Volterra model (hereinafter ‘simple model’); and (ii) ICES‘ (2017) Baltic Sea and ICES‘ (2016) North Sea Ecopath with Ecosim (Christensen and Walters, 2004; Steenbeek et al., 2016) models, both of which were ported into `R` using the ‘`Rpath`’ package (Aydin et al., 2016; Lucey et al., 2020).

Complexity increases with respect to the number of interacting species and trophic links modelled; from 2 species and 2 links for the simple model to 22 and 69 compartments and 82 and 968 links for the Baltic Sea and North Sea models, respectively.

Both the Baltic and North Sea models are used as ‘key-run’ parameterisations peerreviewed by the Working Group on Multispecies Assessment Methods ([WGSAM](https://www.ices.dk/community/groups/pages/wgsam.aspx)). These are the model versions used by ICES to inform management advice.

## 3 Results

Results we present comprise instructions to install the package [note to reviewers: we will update this after depositing the reviewed code on CRAN.], its application to the three benchmark models presented in section 2.6 and performance benchmarks.

### 3.1 Source Code

The package `nash` is available for download at https://github.com/ThomasDelSantoONeill/nash. It can be built directly from source by typing “`install_github(“ThomasDelSantoONeill/nash”)`” in a local `R` session; the `install_github` command is obtained by installing either the “`devtools`” (Wickham et al., 2020) or “`remotes`” (Hester et al., 2020) packages.

### 3.2 The Simple Model

Our two-species simple model is implemented in function `HQLV` in Listing 1. This illustrates an example for the user-supplied `fn` function.

After definition of `HQLV`, the computation of **F**_Nash_ is achieved by calling `nash` for a given set of initial exploitation rates, chosen arbitrarily as 0.2 and 0.3. The output of `nash` is a list with the four elements shown under the ‘`# Results`’ comment in Listing 1. This list informs the user that the `nash` function returned, after 20 function evaluations, *F*_Nash,1_ = 0.41 and *F*_Nash,2_ = 0.44 (dimensions ^1^*/*_Time_) with yield values MSY_1_ = 0.19 and MSY_2_ = 0.22 (dimensions ^Mass^*/*_Time_) respectively.

### 3.3 Ecopath with Ecosim Models

Figure 1 represents snapshots of the Baltic Sea and North Sea food webs, showcasing their distinct complexity. In both networks, non-grey nodes depict the *S* species with commercial value whose harvesting rates we optimised, whilst capturing its effect on the entire food web. To support users employing `Rpath` as their operating model, a built-in function ‘`fn_rpath`’ has been included within the `nash` package to provide the input function `fn` for any `Rpath` model, thus streamlining the interfacing of the two packages. The function `fn_rpath` is capable of transparently handling models in which all or some of the exploited compartments are stage structured (*e*.*g*., all species in the Baltic Sea model and cod, whiting, haddock, saithe and herring in the North Sea model). See Listing 2 for `R` code sample.

**Figure 1:**
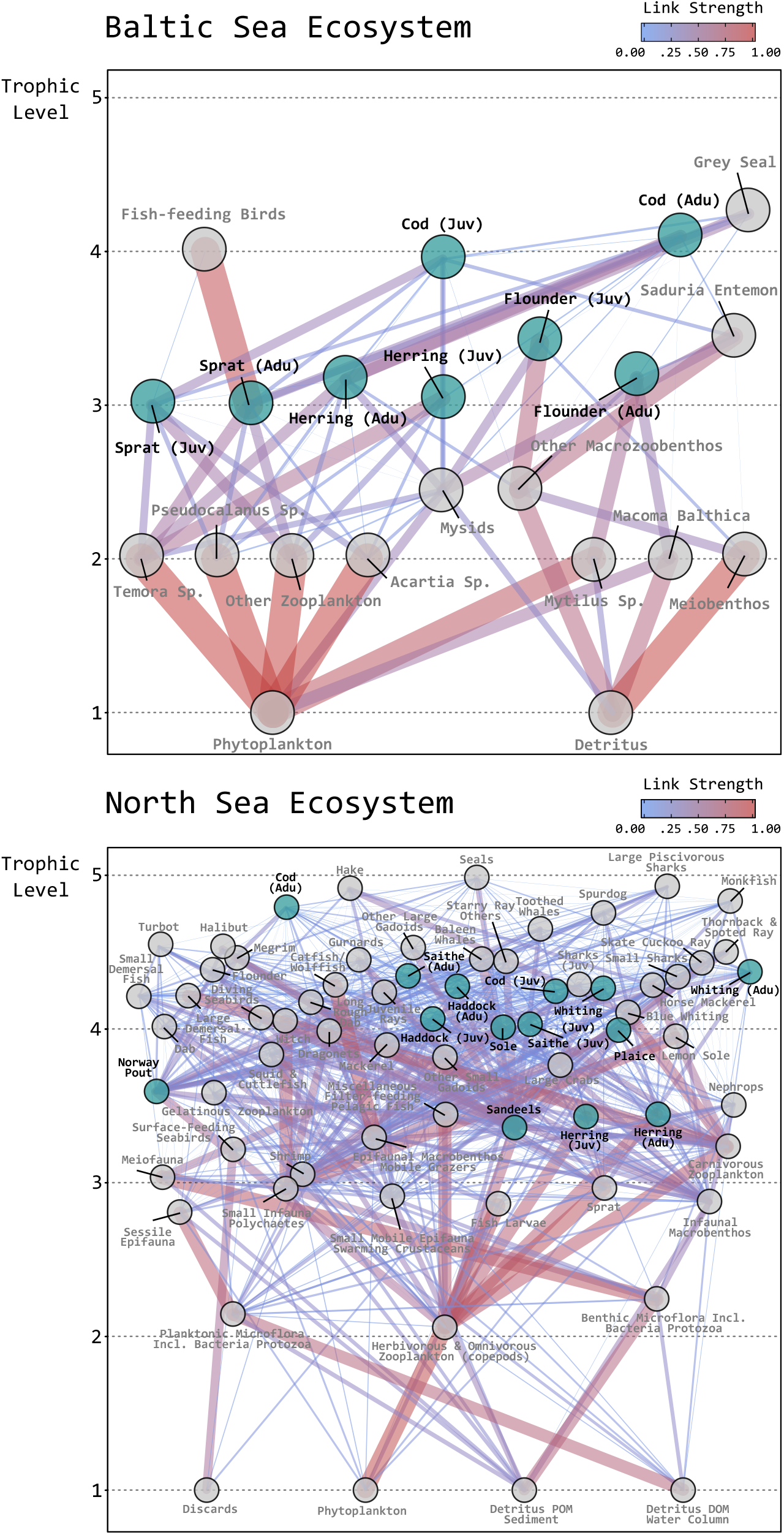
Networks describing the flow of biomass between populations of different species as a result of feeding interactions. Both community food webs are organised by trophic level with non-grey nodes depicting commercial compartments for which `nash` finds optimal harvesting rates taking the entire network as input.

As an easy way to verify NE outputs from the `nash` function, a routine has been included to compute equilibrium yield curves for all *S* exploited species by varying the exploitation rate of one species whilst keeping all others at **F**_Nash_. The user can access this routine by setting `nash`’s logical argument ‘`yield.curves = TRUE`’. Example outputs for both the Baltic Sea and North Sea models are shown, respectively, in Figure 2 and Figure 3. As expected, **F**_Nash_ coincides with the mode of the yield curves in all instances.

**Figure 2:**
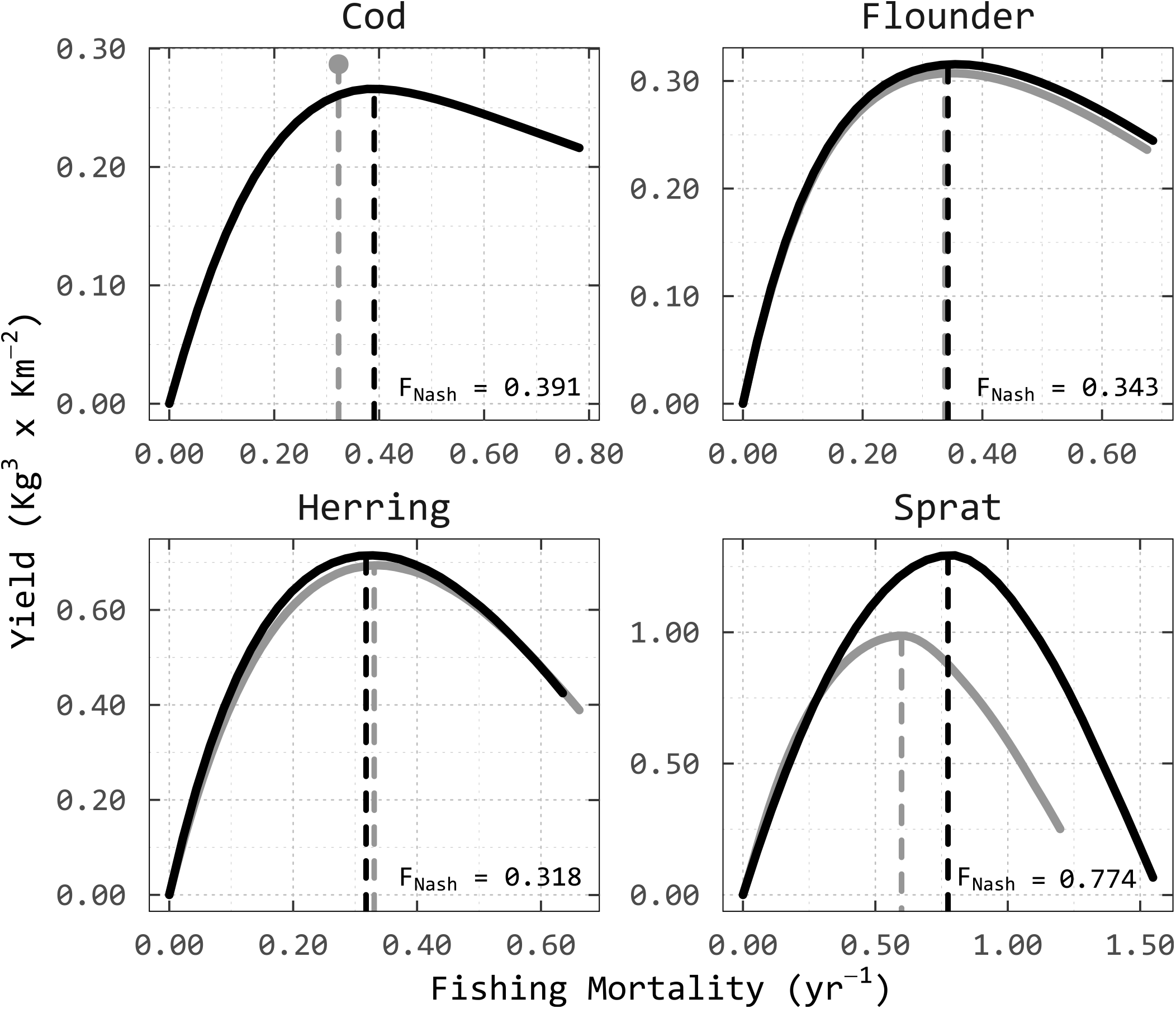
Equilibrium yield curves 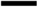 for commercially exploited species in the Baltic Sea ecosystem. The vertical dashed lines 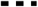 depict the Nash equilibrium harvesting rates found by `nash` for which their values are printed in each frame. Solid grey lines 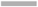 indicate equilibrium yield curves under the constrain of cod remaining 5 *×* 10^4^Kg^3^ above *B*_Nash,Cod_. Likewise, grey vertical dashed lines 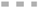 represent a new set of **F**_Nash_ under this conservation constraint. The grey point 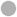 shows the MSY value.

**Figure 3:**
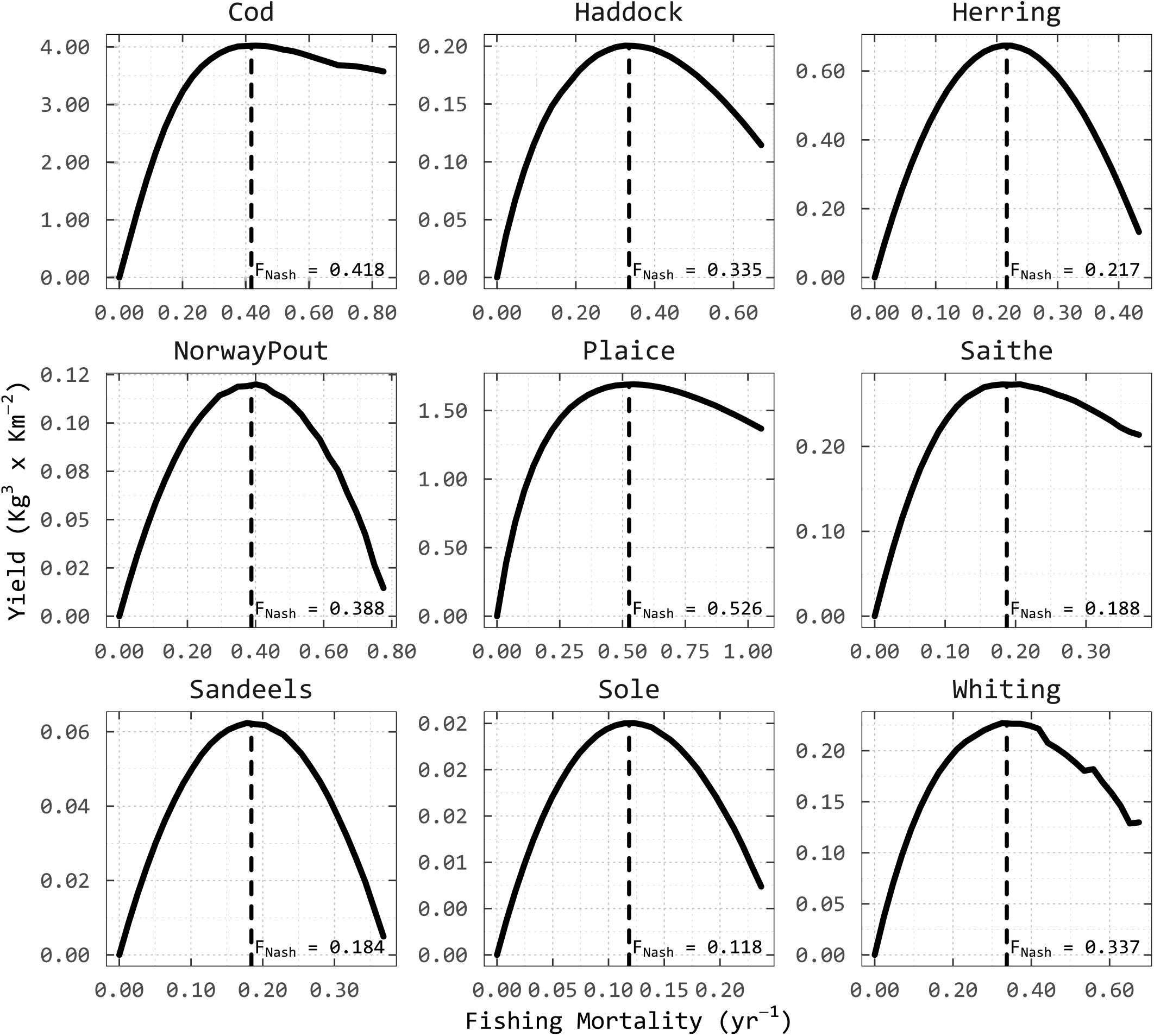
Equilibrium yield curves 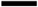 for commercially exploited species in the North Sea ecosystem. The vertical dashed lines 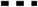 depict the Nash equilibrium harvesting rates found by `nash` for which their values are printed in each frame

In addition, Figure 2 showcases in grey results from adding a conservation constraint for the biomass of cod; which we chose arbitrarily by holding cod 5 *×* 10^4^Kg^3^ above the original *B*_Nash,Cod_ (*i*.*e*. ≈ 21 *×* 10^4^Kg^3^). This results in a new set set of **F**_Nash_ values in which, as expected, the fishing pressure on cod needs to be tapered from 0.391 yr^−1^ to 0.323 yr^−1^. Application of the constraint had little influence on the NE-MSY harvesting rates for herring and flounder, but there is a marginal reduction in their overall yields. For sprat, the conservation constraint on cod led to both lower *F*_Nash_ and lower NE-MSY. By contrast, the constraint on cod biomass generated a surplus of ≈ 5 *×* 10^3^Kg^3^year^−1^ on its yield. Such dependencies amongst fisheries reference points can be evaluated if, and only if, interactions within multispecies complexes are taken into account in models.

### 3.4 nash Benchmarking

The efficiency of the `LV` and the `round-robin` methods was compared by measuring the number of objective function evaluations they require, as a platform-independent metric of performance. For this analysis, we applied both methods across the three tested models with *n* = 29 different initial harvesting rate values. In all cases (see Figure 4), the `LV` method outperformed the `round-robin` method by an approximate factor fifteen in reaching the same convergence threshold (set to a value of 0.01 for this analysis).

**Figure 4:**
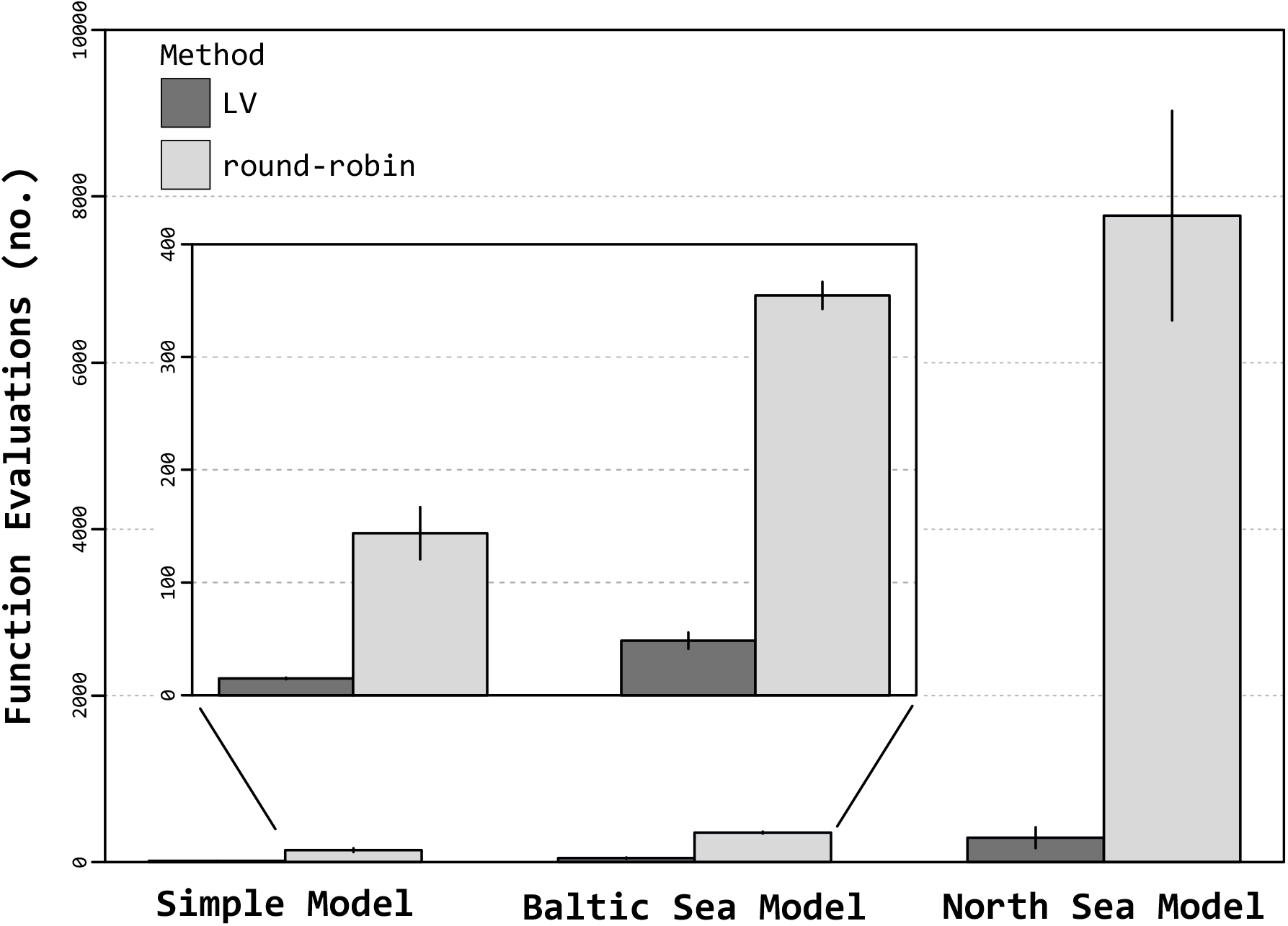
Performance of `nash`’s built-in algorithms measured by the number of objective function calls applied to three models of increasing complexity. Higher bars indicate worse performance with the vertical segments showing one standard deviation about the mean for *n* = 29 initial harvesting rates.

## 4 Discussion

The `R` package `nash` offers an efficient and user friendly route to compute, for a wide range of user-provided multispecies models (*e*.*g*. Plagányi, 2007; Geary et al., 2020), NE-MSY harvesting rates for multiple interacting species simultaneously.

Thorpe et al. (2017) stated that at NE “it is possible to test whether [MSY] have been achieved by varying the fishing mortality on each stock in turn while keeping the others fixed”. This is precisely shown in Figure 2 and Figure 3 for all *S* species and across models, where `nash` correctly computed the NE fishing mortality rates (vertical dashed lines), such that for no stock the long-term yield can be increased by choosing a different fishing mortality. Indeed, the conservation constraint on cod biomass in the Baltic Sea resulted in a decrease of ≈ 65 *×* 10^3^Kg^3^ in overall yield when compared to the overall-unconstrained NE-MSY.

We note that NE-MSY might not necessarily be the optimal solution in all circumstances. For example, NE-MSY does not generally produce the highest sustainble total biomass yield (Farcas and Rossberg, 2016). However, we stress that whenever policies prescribe exploitation of multispecies mixtures at MSY for each interacting species separately, `nash` can help translating this abstract objective into numerical targets.

*Technical interactions*, that is, harvesting more than one species with the same gear concurrently in time and space, are not considered in the sample applications presented herein. Nonetheless, these interactions can be included by modifying `fn` such that it provides the yields of different fleets depending on their efforts.

We briefly consider the `LV` algorithm implemented in `nash` from the computational perspective. Similar to the situation known for the general optimisation problem for an arbitrary univariate objective function over a given domain (Wolpert and Macready, 1997), there appears to be no unique efficient algorithm to find NE in all use cases (Papadimitriou, 2001; Daskalakis et al., 2009). Specific algorithms exploit specific structure in the underlying problems. Our `LV` algorithm exploits that many population dynamical models can, to some extent, be approximated by Lotka-Volterra models, and that for Lotka-Volterra models the NE can be computed analytically using a simple formula. We therefore expect that our algorithm will work well when the problem (*i*.*e*. the ‘game’) is defined by an ecological model, while for other kinds of problems other algorithms will be more suitable (*e*.*g*. the Lemke-Howson algorithm, Lemke and Howson, 1964). For the same reason, we expect that there is generally only a single, pure NE for ecosystem models, while for more generic game-theoretical problems this is not always the case.

The superior performance of `LV` over `round-robin`-like algorithms (Figure 4), for example, is a consequence of the interactions between species, as described by the off-diagonal elements of **G**. For non-interacting species the NE problem reduces to *S* independent maximisation problems, and then `round-robin` will typically be more efficient because it solves the *S* problems independently. At first sight, differences in computation time between the algorithms might appear negligible compared to the time and effort required to construct finely parameterised complex models. However, when performing sensitivity analyses, management strategy evaluations (MSE), or when exploring model variants where many NE need to be computed, `nash`’s `LV` routine can bring substantial improvements in computation times.

The `nash` package tool was built with the intention to streamline the computation of NE where the ‘game’ is defined by an ecological model. As a result, its application is particularly well-suited for communities of interacting species.

We believe that `nash` has the potential to overcome current barriers to full implementation of rational ecosystem-based management policies (*e*.*g*. the MSFD or, more recently, UK’s Joint Fisheries Statement), thus ultimately contributing towards the realisation of the United Nations’ post-2015 global sustainability targets set forth in A/RES/70/1 with the adoption of “Transforming our world: the 2030 Agenda for Sustainable Development” (available at https://digitallibrary.un.org/record/3923923?ln=en).

## Acknowledgements

The development of `nash` is founded by UK’s Research and Innovation (UKRI) Biotechnology and Biological Sciences Research Council (BBSRC; project identifier BB/M009513/1 number 2243988) and the Centre for Environment, Fisheries and Aquaculture Science (CEFAS; project identifier DP902I). We would like to acknowledge the feedback provided by Dr. Sean M. Lucey (NOAA’s Northeast Fisheries Science Center) and Dr. Kerym Aydin (NOAA’s Alaska Fisheries Science Centre) with regards to `Rpath`’s support.

## Authors’ Contributions

AGR conceived of the study. TJDSO and AGR designed and scripted the source package. TJDSO developed the models, performed simulations, analysed the data and drafted the manuscript. All authors contributed critically to the drafts and gave final approval for publication. The authors declare no competing interests.

## Algorithm 1: ‘LV’ method

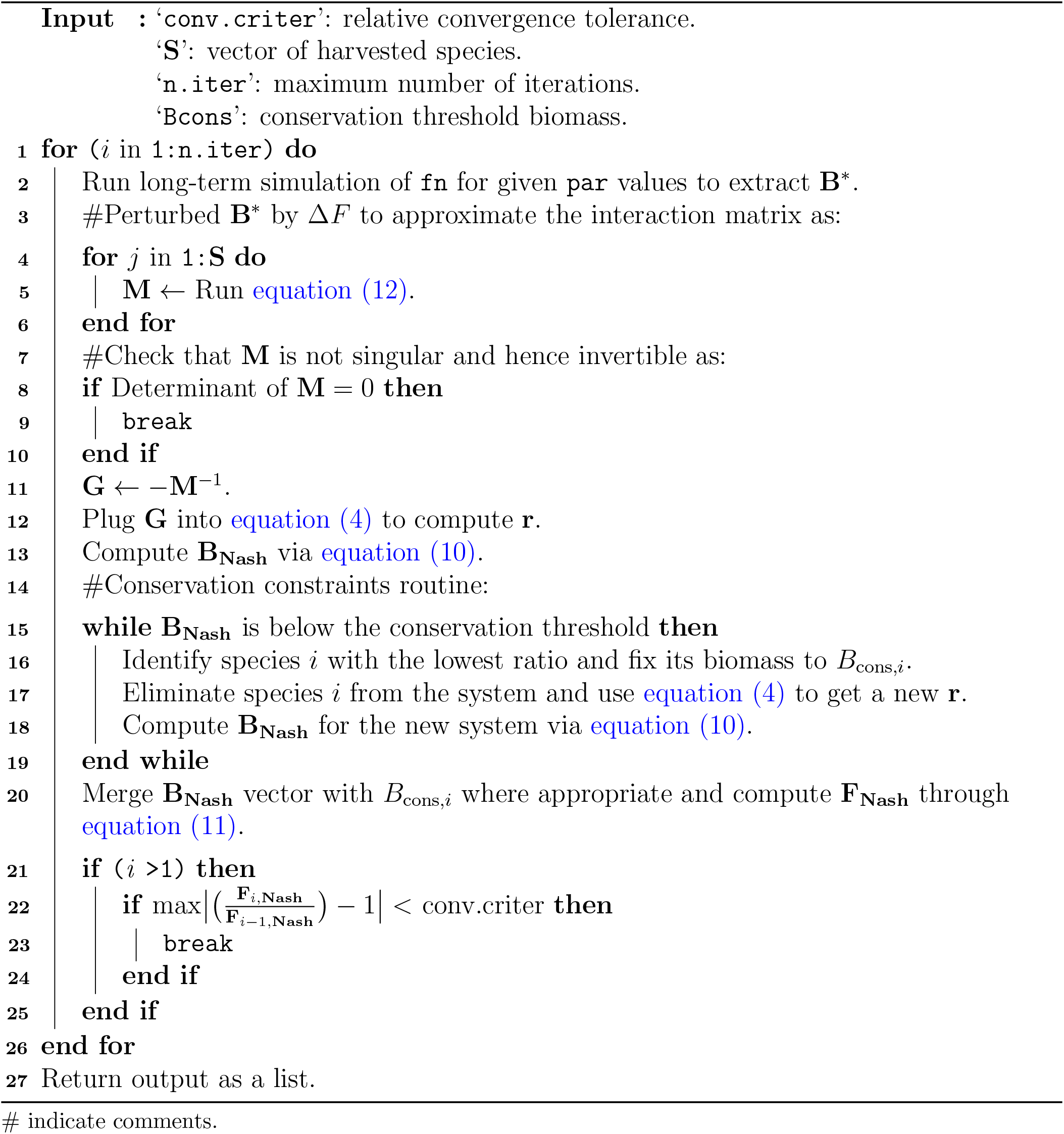

## Algorithm 2: ‘round-robin’ method

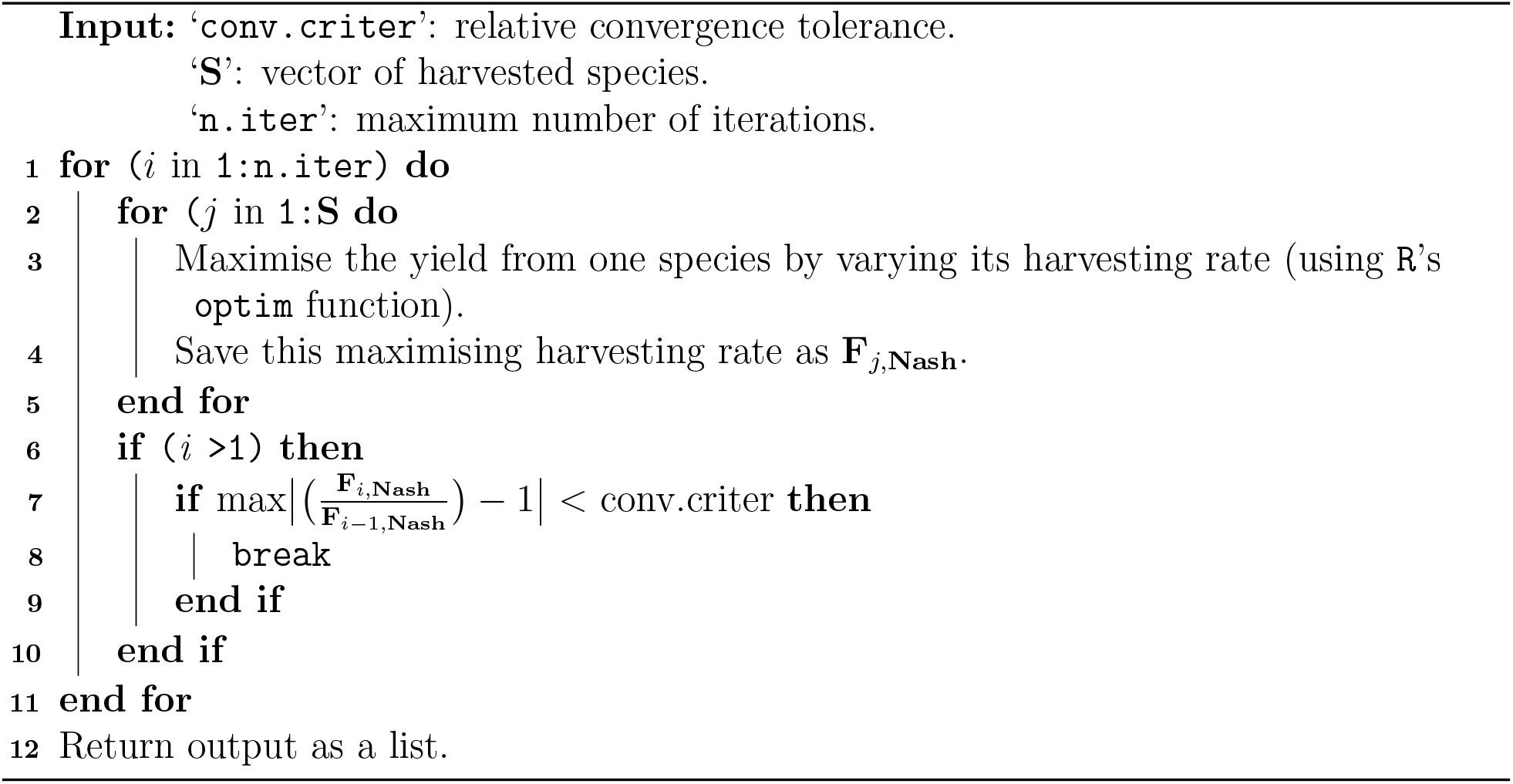

**Listing 1:**
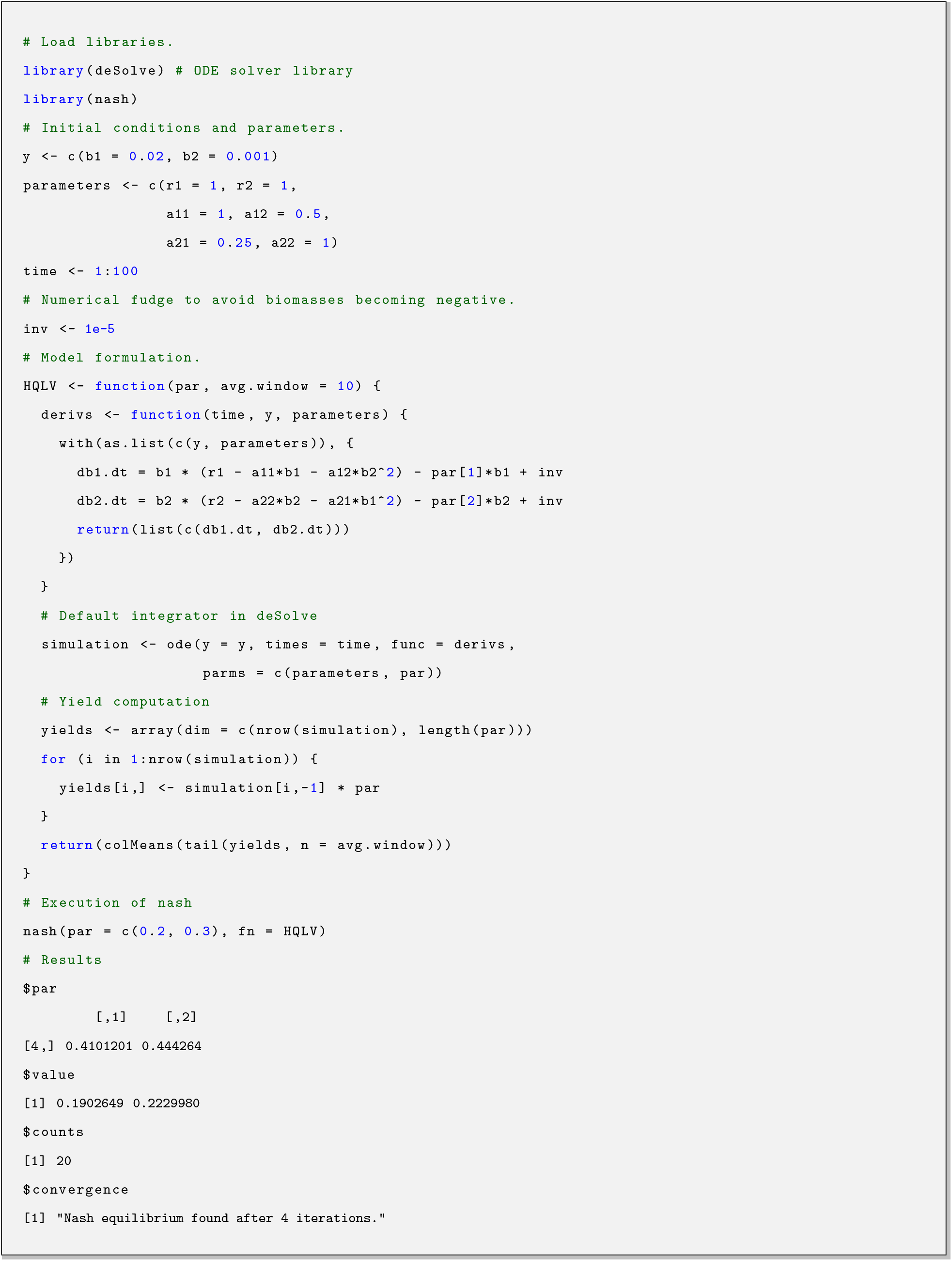
`R` code illustrating the application of `nash` to the simple model. See Soetaert et al., 2010 for `deSolve` details.

**Listing 2:**
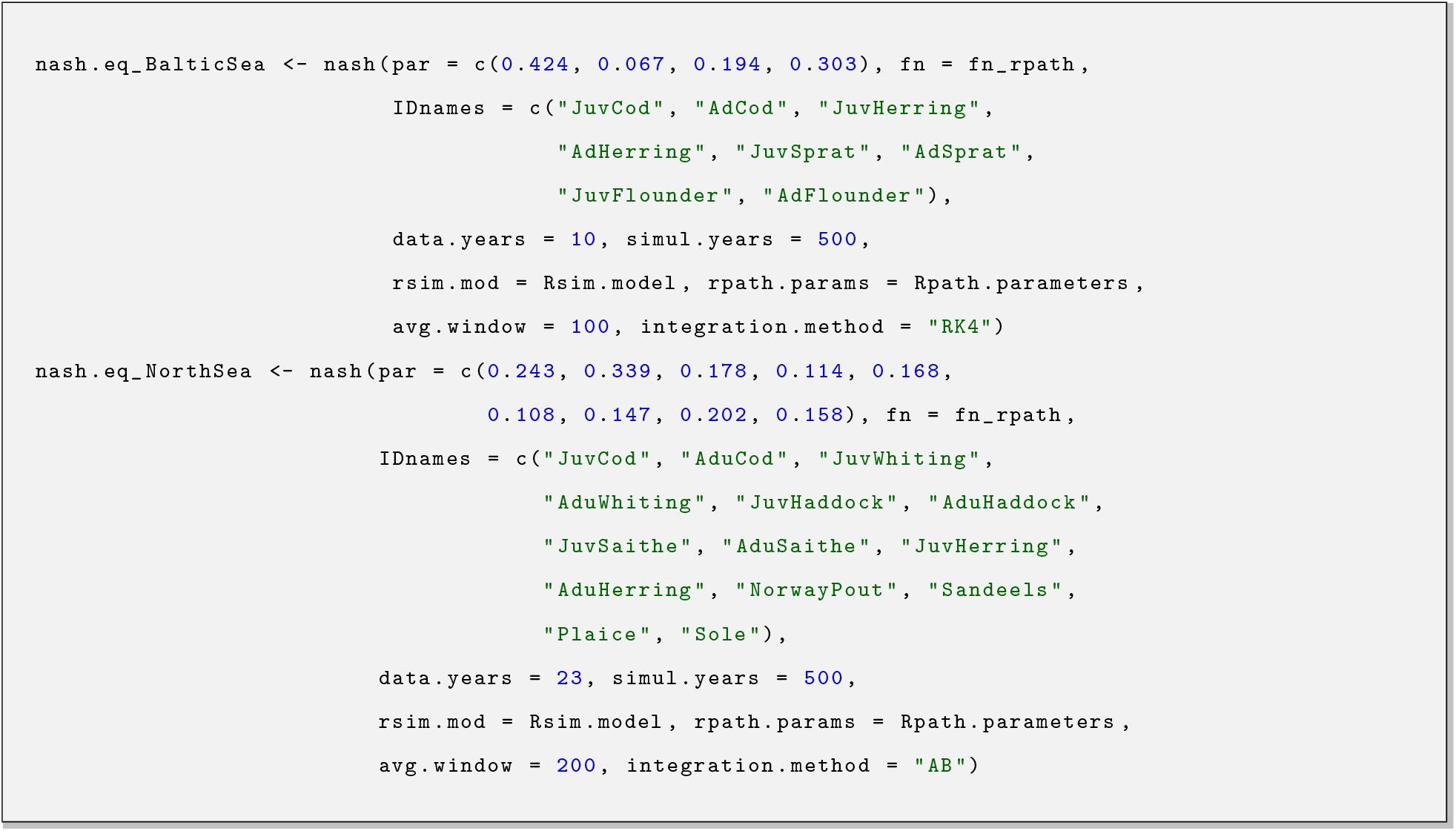
Application of `nash` to the Baltic Sea and North Sea `Rpath` models. In both calls, all the parameters after `fn` are passed to `fn_rpath`. ‘`Rsim.model`’ and ‘`Rpath.parameters`’ represent `R` objects within which the scenario for dynamic simulations and parameters for mass balance are stored. A thorough documentation file can be accessed running `?fn_rpath` command.

## Footnotes

1 Currently being superseded by the Nature’s Contribution to People approach (Díaz et al., 2018) as a result of internal debates within the Intergovernmental Science-Policy Platform on Biodiversity and Ecosystem Services (IPBES) addressing the shortcomings of utilising an ecosystem services framework to promote environmental protection (however see Muradian and Gómez-Baggethun, 2021).

2 By stock of fish we mean a self-contained geographical grouping of fish species that share a common reproductive process.

3 The term Sustained Yield (SY) is conceptually synonymous with MSY. SY chronologically precedes MSY with its first occurrence attributed to von Carlowitz (1713) with reference to forestry.

4 Opened for signature on December 10, 1982 and, in accordance with article 308(1), entered into force on November 16, 1994. As of January 25, 2023, 168 states and the European Union are parties to the Convention (available at https://www.un.org/depts/los/conventionagreements/convention_overview_convention.htm).

5 Pub. L. 94–265, Apr. 13, 1976, 90 Stat. 331 as amended by the Magnuson-Stevens Fishery Conservation and Management Reauthorization Act of 2006 Pub. L. 109-479, Jan. 12, 2007, 120 Stat. 3575 (for complete classification see 16 U.S.C. 1801 *et seq* at https://www.govinfo.gov/content/pkg/USCODE-2017-title16/html/USCODE-2017-title16-chap38.htm).

6 Regulation (EU) No 1380/2013 of the European Parliament and of the Council of 11 December 2013 on the Common Fisheries Policy, amending Council Regulations (EC) No 1954/2003 and (EC) No 1224/2009 and repealing Council Regulations (EC) No 2371/2002 and (EC) No 639/2004 and Council Decision 2004/585/EC (OJ L 354 28.12.2013, pp. 22-61 as amended by OJ L 354 28.12.2013, pp. 86-89; OJ L 133 29.5.2015, pp. 1-20; OJ L 302 17.11.2017, pp. 1-2; and OJ L 198 25.7.2019, pp 105-201). Available *mutatis mutandis* (*m*.*m*) at http://data.europa.eu/eli/reg/2013/1380/2019-08-14.

7 A Bill to make provision in relation to fisheries, fishing, aquaculture and marine conservation; to make provision about the functions of the Marine Management Organisation; and for connected purposes. Formally introduced in the House of Lords on January 29, 2020 and passed to the House of Commons for its first reading on July 2, 2020. The Bill became an Act of Parliament (law) after receiving Royal Assent on November 23, 2020 (available at https://www.legislation.gov.uk/ukpga/2020/22/contents).

8 Directive 2008/56/EC of the European Parliament and of the Council of 17 June 2008 establishing a framework for community action in the field of marine environmental policy (Marine Strategy Framework Directive) (Text with EEA relevance). OJ L 164, 25.6.2008, p. 19–40 as amended by OJ L 125, 18.5.2017, p. 27–33; available *m*.*m*. at https://eur-lex.europa.eu/eli/dir/2008/56/2017-06-07.

9 Regulation (EU) 2016/1139 of the European Parliament and of the Council of 6 July 2016 establishing a multiannual plan for the stocks of cod, herring and sprat in the Baltic Sea and the fisheries exploiting those stocks, amending Council Regulation (EC) No 2187/2005 and repealing Council Regulation (EC) No 1098/2007 (OJ L 191, 15.7.2016, p. 1–15 as amended by OJ L 179, 16.7.2018, p. 76–77; OJ L 83, 25.3.2019, p. 1–17; OJ L 198, 25.7.2019, p. 105–201; and OJ L 400, 30.11.2020, p. 1–6). Available *m*.*m*. at http://data.europa.eu/eli/reg/2016/1139/2020-12-01. Regulation (EU) 2018/973 of the European Parliament and of the Council of 4 July 2018 establishing a multiannual plan for demersal stocks in the North Sea and the fisheries exploiting those stocks, specifying details of the implementation of the landing obligation in the North Sea and repealing Council Regulations (EC) No 676/2007 and (EC) No 1342/2008 (OJ L 179, 16.7.2018, p. 1–13 as amended by OJ L 83, 25.3.2019, p. 1–17; and OJ L 198, 25.7.2019, p. 105–201). Available *m*.*m*. at http://data.europa.eu/eli/reg/2018/973/2019-08-14.

